# To Predict is to Design: Unlocking Generative Capabilities in All-Atom Structure Predictors via Geometric Score Distillation

**DOI:** 10.64898/2025.12.23.696242

**Authors:** Yuxuan Li, Yeyu Su, Yanbo Jing, Tao Liu

## Abstract

Current protein binder design largely relies on a decoupled paradigm: generating backbones via unconditioned diffusion followed by sequence filling or refilling with inverse folding models. This separation prevents the design process from accessing the holistic validation metrics of structure predictors during generation, wasting rich physical priors. While recent works like BindCraft have successfully inverted AlphaFold2 for protein design^1^, extending this inversion to state-of-the-art all-atom diffusion predictors (e.g., AlphaFold3, Boltz-2) remains a formidable challenge, particularly for modalities requiring non-standard residues such as cyclic peptides. In this work, we present DREAM (<u>D</u>ifferentiable <u>R</u>efinement via <u>E</u>nergy-<u>A</u>nchored <u>M</u>anifolds), a model-agnostic framework that turns the passive predictive trajectory of diffusion models into an active, lucid design process. DREAM repurposes Boltz-2^2^—a leading open-source all-atom predictor—via Geometric Score Distillation (GSD), a technique enabling explicit gradient-based optimization directly through the frozen diffusion network. Unlike previous methods constrained by standard amino acids, DREAM directly unlocks the model’s latent chemical vocabulary, allowing gradients to autonomously select the optimal building blocks up to 55 residue types (including D-amino acids and post-translational modifications) to minimize energy. We demonstrate this capability by designing cyclic peptide binders for diverse targets, including PD-L1, B7-H3, and the human μ-Opioid Receptor (hMOR). Our results suggest that the programmable design of chemically complex modalities is not a distant goal, but a latent capability of current all-atom predictors, waiting to be inverted. Ultimately, to predict is to design.

## Introduction

The revolution in protein structure prediction, marked by AlphaFold2^3^ and recently culminated in all-atom diffusion models like AlphaFold3^4^, Boltz-2^2^, and Protenix^5^, has fundamentally solved the static folding problem for a vast majority of biomolecules. These models have internalized a profound understanding of biophysics, capturing the nuances of side-chain packing, ligand interactions, and even post-translational modifications. However, a significant dichotomy remains: while we possess oracles capable of predicting complex structures with high fidelity, harnessing these models to generate novel functional proteins—particularly those with non-standard chemistries—remains a formidable challenge.

Current state-of-the-art design methods predominantly follow a “generate-then-validate” or decoupled paradigm. Specialized generative models like RFdiffusion1, 2 and 3 construct backbones in a latent space, relying on external inverse folding tools like ProteinMPNN^6^ or LigandMPNN^7^ to fill or refill in sequences^8–10^. While effective for standard proteins, this approach is fundamentally limited by the training data and fixed vocabulary of the generator. Extending such models to include non-standard residues (e.g., D-amino acids) or complex modifications requires expensive retraining. Alternatively, “hallucination” methods attempt to invert prediction models^1^. While successful with AlphaFold2, applying this inversion to stochastic diffusion models (AF3/Boltz-2) is non-trivial. Alternative strategies, including recent efforts like BoltzDesign, attempt to bypass the diffusion bottleneck by restricting optimization to auxiliary outputs such as distograms or confidence heads. While this approach enables gradient backpropagation, it optimizes probabilistic proxies rather than the explicit 3D coordinates, thereby discarding the precise atomic-level geometric information (e.g., side-chain packing) that only emerges after the full diffusion decoding^11^. Recent attempts relying on iterative forward passes are fundamentally limited to stochastic sampling of the model’s manifold^12,13^. Such approaches tend to converge toward the model’s highest-likelihood (and thus most typical) priors, limiting their ability to actively navigate toward low-probability, high-utility regions required for specific functional goals.

We believe that the long-standing desideratum of programmable molecular design is not to train new models, but to invert the powerful predictors we already possess. To predict is to design. A fully trained all-atom predictor like Boltz-2 contains a latent generative capability far richer than any specialized model, covering a chemical vocabulary of 55 residue types and complex geometric priors. The challenge lies in extracting this capability: how do we turn the stochastic, noise-driven diffusion process into a deterministic, goal-directed optimization?

In this work, we introduce DREAM (Differentiable Refinement via Energy-Anchored Manifolds), a model-agnostic framework that unlocks the generative potential of frozen all-atom predictors (Scheme 1). Unlike standard score distillation sampling (SDS) used in image generation^14^, we propose Geometric Score Distillation (GSD) to circumvent the long-standing instability and vanishing gradients inherent in backpropagating through hundreds of diffusion sampling steps. By treating the single-step denoising operation as a differentiable projector, we backpropagate explicit geometric gradients—such as contact constraints or radius of gyration— directly through the frozen diffusion network to the sequence logits. This transforms the predictor from a passive oracle into an active design engine, allowing us to “steer” the diffusion trajectory toward functional high-energy states that random sampling would hardly visit.

Importantly, DREAM eliminates the reliance on external sequence design tools. By optimizing sequence logits directly against the predictor’s all-atom energy landscape, our framework autonomously selects from an extended vocabulary of 55 residue types—including D-amino acids and phosphorylated residues—to resolve potential steric clashes or improve binding probability. We demonstrate the versatility of DREAM by designing cyclic peptide binders for diverse targets, including PD-L1, B7H3, and the human μ-Opioid Receptor (hMOR). Our results show that by simply providing a gradient compass, we can navigate the vast, rugged landscape of all-atom diffusion models to discover novel, chemically complex molecules.

**Scheme 1:**
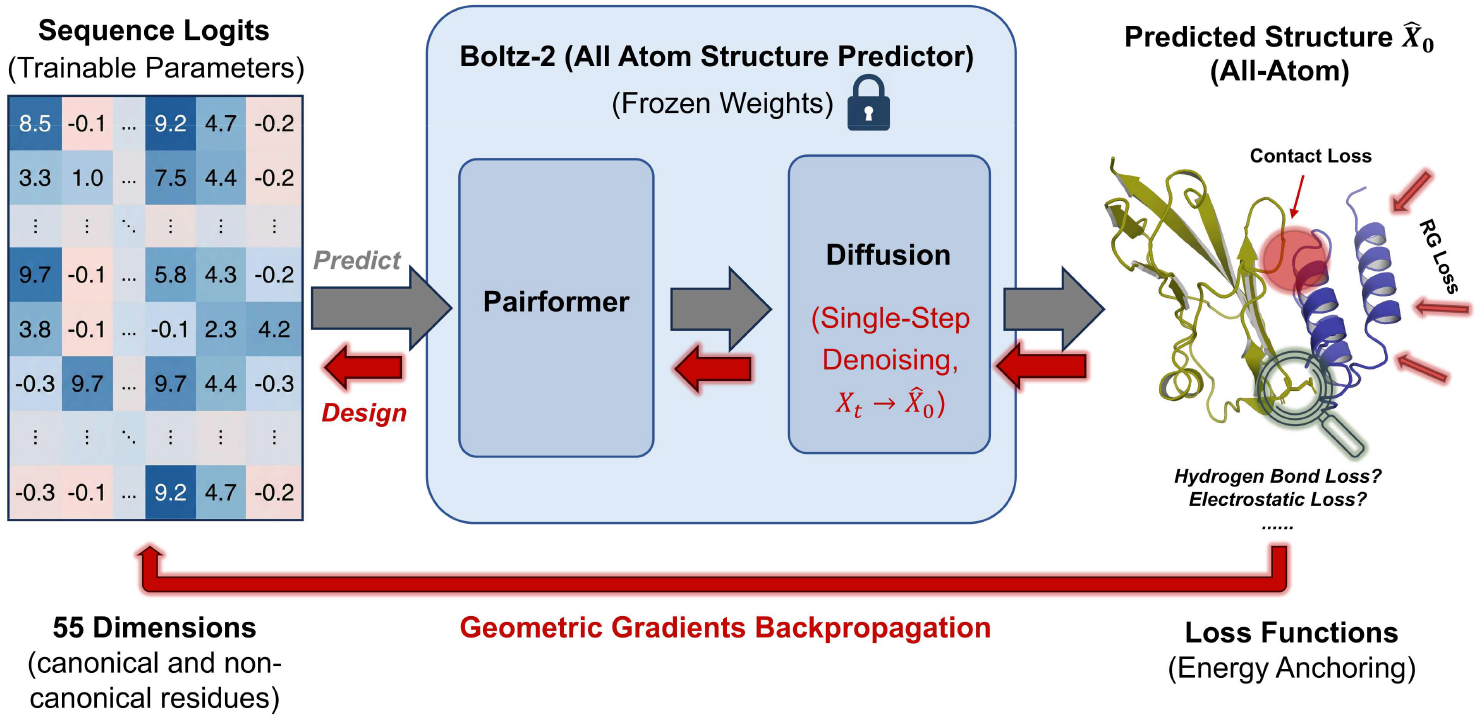
The DREAM Framework for Differentiable Design. Starting from 55-dimensional sequence logits (supporting non-standard residues), the model generates an immediate structural estimate 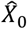 via single-step denoising. Explicit geometric losses (e.g., Contact, RG) are calculated on this structure, and gradients are backpropagated (red arrow) through the frozen network to actively update the sequence. We term this process Geometric Score Distillation (GSD).

## The DREAM Framework

### 1. Energy-Anchored Manifolds: The Single-Step Denoising Proxy

We present DREAM (Differentiable Refinement via Energy-Anchored Manifolds), a general framework designed to invert state-of-the-art all-atom structure prediction models. While recent proprietary models like AlphaFold3^4^ have demonstrated unprecedented accuracy, its closed nature limits its broad utility for de novo design. DREAM is architected to be model-agnostic. In this work, we implement DREAM on Boltz-2^2^, one of the current state-of-the-art open-source all-atom predictor, utilizing it as a differentiable engine to map sequence logits to structural geometries. A central challenge in diffusion-based optimization is the unreachable computational cost and the instability arising from exploding/vanishing gradients of differentiating through long sampling trajectories^15^ (e.g., 200 steps). Here we propose a radical simplification: calculating geometric losses on the single-step denoised estimate 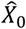 derived from a noisy state *X*_t_. This approach was grounded in Tweedie’s Formula, a foundational result in diffusion theory^16–18^. Tweedie’s Formula states that the single-step denoised output 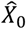 is theoretically equivalent to the posterior expectation (or Minimum Mean Squared Error estimator) of the clean data given the noise:

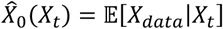

This mathematical property implies that 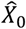 is not merely a “blurry sample,” but the statistical center of mass of the valid structural manifold compatible with the current noise level^18^. At high noise levels (High σ), 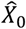 captures the global topology and compactness. At low noise levels (Low σ), 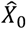 converges to precise atomic details.

Therefore, optimizing a loss function on this expectation 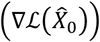 provides a statistically valid gradient direction that steers the sequence distribution towards the desired geometric manifold, effectively bypassing the need for expensive full trajectory unrolling. While high-noise regimes (σ > 100) excel at shaping global topology by targeting the structural ‘center of mass’, accessing atomic-level physicochemical features (e.g., specific hydrogen bonds) requires mitigating this averaging effect. This necessitates operating in low-noise regimes, where the input *X*_t_ must lie near the valid data distribution. To bridge this gap, we employ an anchor strategy: instead of starting from random noise, we initialize *X*_t_ by perturbing a generated structure (the ‘anchor’) derived from the current sequence. This is the literal manifestation of the ‘Energy-Anchored’ concept in our DREAM acronym. By anchoring the gradient calculation to a valid, low-energy conformation rather than the void of random noise, we ensure that the optimization remains strictly on-manifold^19^. This strategy effectively combines the global steering capability of high-sigma diffusion with the atomic precision of low-sigma refinement, a capability that will be further expanded in future iterations.

### 2. Geometric Score Distillation: Differentiable Steering through the Frozen Predictor

To navigate this manifold, we introduce <u>G</u>eometric <u>S</u>core <u>D</u>istillation (GSD). While inspired by Score Distillation Sampling (SDS) used in image generation^14^ (e.g., DreamFusion), our approach diverges fundamentally in how gradients are computed to suit the precise nature of molecular design. Standard SDS typically optimizes a probabilistic objective and bypasses the computationally expensive Jacobian of the denoising network for stability^14^. Our GSD, conversely, aims to satisfy explicit Euclidean constraints (e.g., contact maps, radius of gyration). We contend that the structural logic encoded within the diffusion network—the mapping from sequence to geometry—is crucial for satisfying these constraints. Therefore, we explicitly backpropagate through the frozen diffusion network, retaining the Jacobian term. The gradient update for our sequence parameters *θ* is derived via the chain rule:

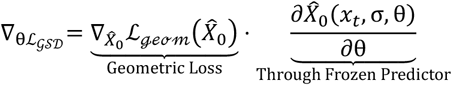

Here, 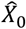 acts as a differentiable bridge. By retaining the Jacobian term 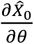, DREAM allows the geometric “desire” (e.g., “close this loop”) to be translated directly into specific chemical mutations (e.g., “mutate Gly to Cys”) via the predictor’s learned physics. We term this “Geometric Score Distillation” because we are distilling the geometric priors encoded in the score function directly into the design space.

## Results

### 1. DREAM-boltz2 optimize structure and sequence simultaneously

To demonstrate the capability of DREAM-boltz2, we first applied the framework to the de novo design of a cyclic peptide binder. The optimization workflow (Figure 1a) initiates from high-noise states (σ = 500), performing a single-step diffusion to estimate the structure 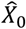. Gradients are then backpropagated from three minimalistic loss functions: an ipTM loss and pLDDT loss derived from the model’s confidence heads, and a hotspot loss requiring contact with a target residue. Crucially, we restricted the design space to the 20 canonical amino acids without imposing any explicit sequence constraints (e.g., diversity or hydrophobicity penalties), relying solely on the model’s inherent physical intuition to orchestrate the sequence composition.

**Figure 1:**
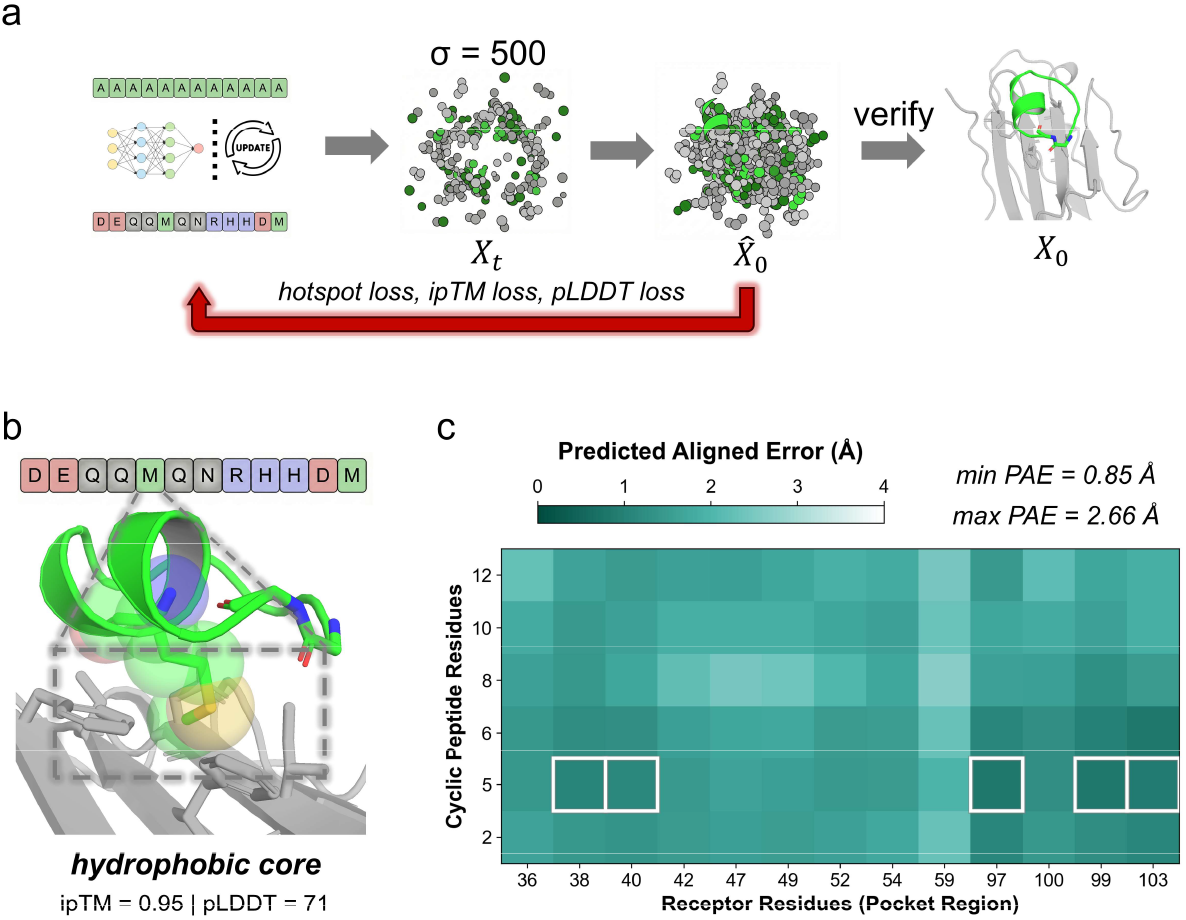
Simultaneous Sequence-Structure Co-Optimization of DREAM-boltz2. (a) The gradient-driven design workflow. Optimization initiates from high-noise states (sigma = 500), utilizing single-step denoising to estimate the structural expectation*X*0_hat. Geometric gradients from minimal objectives (hotspot, ipTM, pLDDT) are backpropagated through the frozen predictor to steer sequence evolution. (b) Representative cyclic peptide design targeting the flat surface of B7-H3. The optimizer spontaneously selected Methionine (M) residues to bury a shallow surface depression, creating a precise hydrophobic core (highlighted) to anchor the binder. (c) Interface Predicted Aligned Error (PAE) matrix. The deep teal regions (highlighted boxes) indicate sub-angstrom confidence at the core interface (min PAE = 0.85 Angstroms), confirming the model’s certainty in the rigid, high-affinity complex formation.

We selected B7-H3, a clinically valuable immune checkpoint target^20^, for this validation. Structurally, B7-H3 is characterized by Immunoglobulin-like domains presenting a flat surface that lacks deep pockets, posing a challenge for binder design^20^. We initialized a 12-residue cyclic peptide and allowed DREAM to simultaneously evolve both its topology and sequence. The optimization trajectory successfully identified and exploited key surface features of the target. As shown in Figure 1b, the optimizer identified a shallow hydrophobic depression on the flat B7-H3 surface. To fill this, it spontaneously selected a Methionine (M) residue, positioning its bulky side chain to bury the hydrophobic surface area, thereby creating a tight hydrophobic core. Remarkably, despite the absence of sequence constraints, the resulting peptide exhibited a “healthy” solubility profile: aside from the two hydrophobic residues (M) forming the core, the remaining residues were hydrophilic. This suggests the model inherently balances binding affinity with solvent exposure.

Quantitative analysis further validates the design quality. The Predicted Aligned Error (PAE) matrix (Figure 1c) shows consistently low error (< 1 Å) at the specific hydrophobic core interface, with the remaining interface regions below 2.7 Å. This confirms that even at high noise levels (σ = 500), the gradients derived via GSD provides a valid and precise steering signal. The model successfully recognized the receptor’s surface energetics and performed accurate “shape and chemical matching” in a single co-optimization loop, validating our hypothesis that the frozen predictor possesses latent, actionable physicochemical knowledge.

### 2. Unsupervised Emergence of Complex Medicinal Chemistry Strategies from DREAM-boltz2

To transcend the limitations of canonical protein design, we expanded the DREAM optimization manifold to encompass the full chemical vocabulary of Boltz-2. By unlocking the logits to 55 dimensions—integrating 20 canonical amino acids with 35 non-canonical types (Figure 2a)—we enabled the model to navigate a vastly expanded fitness landscape, selecting building blocks based purely on intrinsic biophysical necessity. The framework first manifested an emergent understanding of chiral geometry during the design of a cyclic binder for B7-H3. The optimizer spontaneously incorporated a D-Tryptophan (D-Trp) into a constrained segment of the macrocycle. Structural interrogation (Figure 2b) reveals this was a masterstroke of geometric precision: substituting D-Trp with its L-enantiomer triggers a severe steric clash between the bulky indole side chain and the backbone carbonyl of the i+3 residue. The model did not merely “suggest” a mutation; it leveraged non-standard chirality to unlock a stable, low-energy conformation that is physically forbidden to canonical L-peptides, demonstrating a deep, latent grasp of stereo-chemical constraints^21^.

**Figure 2:**
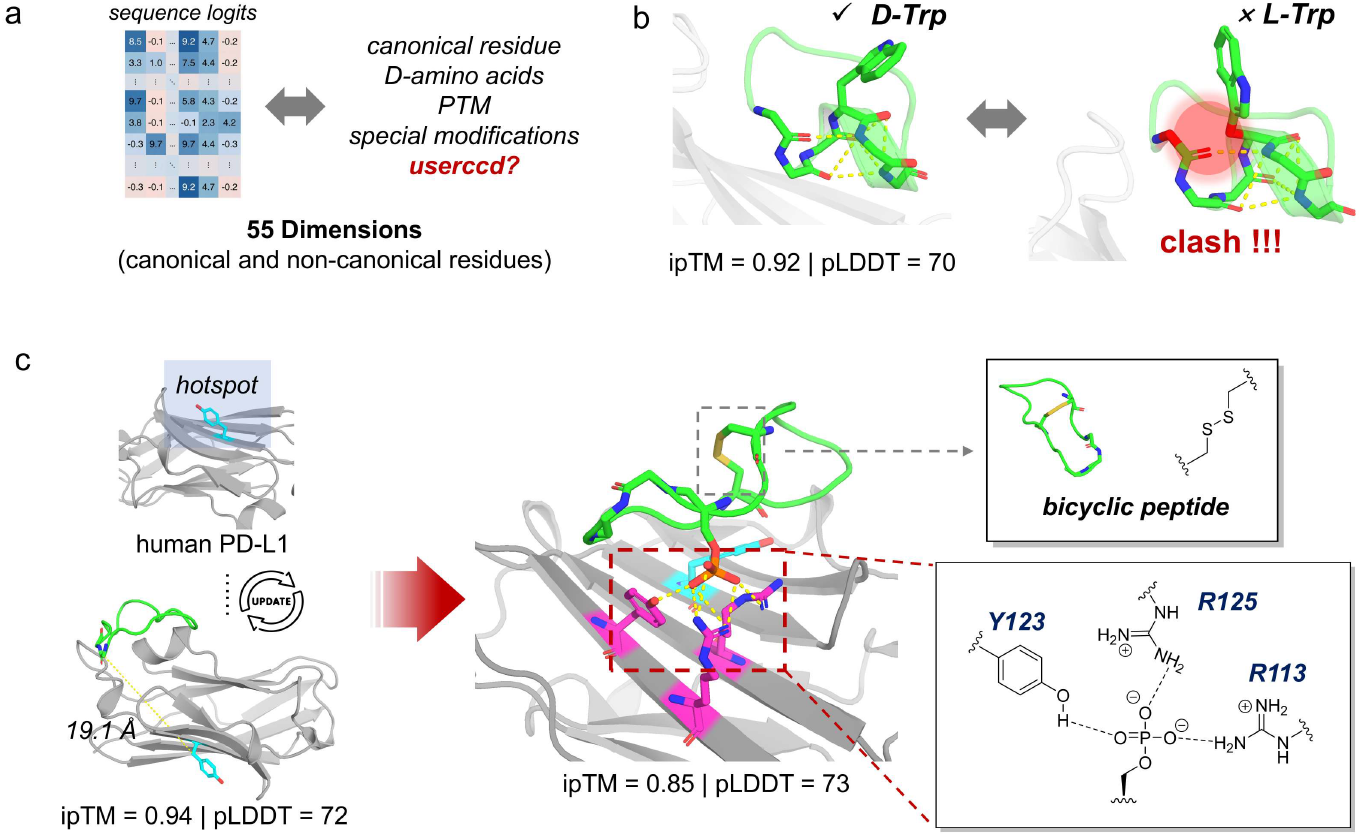
Unlocking Latent Chemical Vocabulary and Emergent Structural Complexity of DREAM-boltz2. (a) DREAM-boltz2 expands the optimization space to a 55-dimensional manifold, integrating canonical amino acids with non-canonical types (e.g., D-amino acids, PTMs) to enable physics-driven selection. (b) Geometric Emergence: In the B7-H3 binder design, the model autonomously incorporated a D-Tryptophan (D-Trp). Substituting this with the L-enantiomer triggers a severe steric clash (red glow) with the backbone, demonstrating the model’s capacity to utilize non-standard chirality to access geometries forbidden to canonical peptides. (c) Physical and Topological Emergence: Design targeting the flat, hydrophilic interface of PD-L1. Although initialized as a simple loop, the model spontaneously evolved a figure-8 bicyclic scaffold (inset) via a de novo disulfide bond to minimize conformational entropy. Simultaneously, the optimizer deployed a Phosphoserine (SEP) residue to anchor a dense polar triad (Y123, R125, R113). This forms a sub-angstrom electrostatic clamp—a sophisticated salt-bridge network that standard residues could not satisfy—effectively sealing the interface.

The model’s “design logic” became even more striking when confronted with the challenging, flat and hydrophilic interface of human PD-L1^22^. Starting from a featureless 15-mer monocyclic loop, the gradient descent process performed a spontaneous topological evolution: it introduced two cysteine residues that self-organized into a rigid figure-8 bicyclic scaffold (Figure 2c). Remarkably, this transition occurred without any pre-defined templates or bias toward disulfide formation. This suggests that the optimization landscape naturally funnels toward entropy reduction; the model “discovered” that partitioning the large flexible loop into a bicyclic constraint was the most efficient way to maximize binding confidence (pLDDT) and minimize the conformational penalty on the flat target surface^23^.

Simultaneously, the model resolved the complex interface chemistry by deploying a Phosphoserine (SEP) to anchor a triad of polar residues (Y123, R125, R113). While standard residues like Asp or Glu provide negative charge, they lack the specific charge density and tetrahedral geometry required to satisfy this dense polar cluster. The model’s autonomous selection of SEP functions as a sub-angstrom electrostatic clamp, forming a sophisticated salt-bridge network that effectively “seals” the interface (Figure 2c). This emergent physical logic— utilizing covalent bicyclic constraints and high-density electrostatic anchors—confirms that DREAM is not merely reciting its training data, but is acting as a biophysics-aware engine capable of deriving de novo chemical solutions to intractable design problems.

### 3. DREAM-boltz2 shows potential in multiple targets

To demonstrate the translational potential of DREAM-boltz2, we challenged the framework with two other therapeutic targets: the “undruggable” oncogene KRAS-G12D and the human μ-Opioid Receptor (hMOR). Targeting the shallow, featureless surface of KRAS-G12D is difficult, as it offers few pocket-like anchors for binding^24^. We restricted the design space to a compact 10-residue cyclic peptide. This constraint demands a low entropic penalty upon binding, which the DREAM optimizer addressed by orchestrating a sophisticated non-canonical ensemble. The model strategically utilized D-Proline and D-Phenylalanine to induce a sharp, rigid reverse turn, allowing the 10-mer macrocycle to execute a “pincer maneuver” that probes deep into the hydrophobic cleft (Figure 3a). Another evidence of emergent chemical logic was the spontaneous selection of α-aminoisobutyric acid (Aib). By leveraging the gem-dimethyl effect of Aib, the model performed “steric steering”—physically pushing against a proximal Methionine to lock the short peptide into its bioactive conformation. This demonstrates that within the spatial constraints of a 10-mer, DREAM-boltz2 can utilize non-standard building blocks as de novo structural scaffolding to maximize geometric and chemical complementarity on shallow, traditionally intractable interfaces.

**Figure 3:**
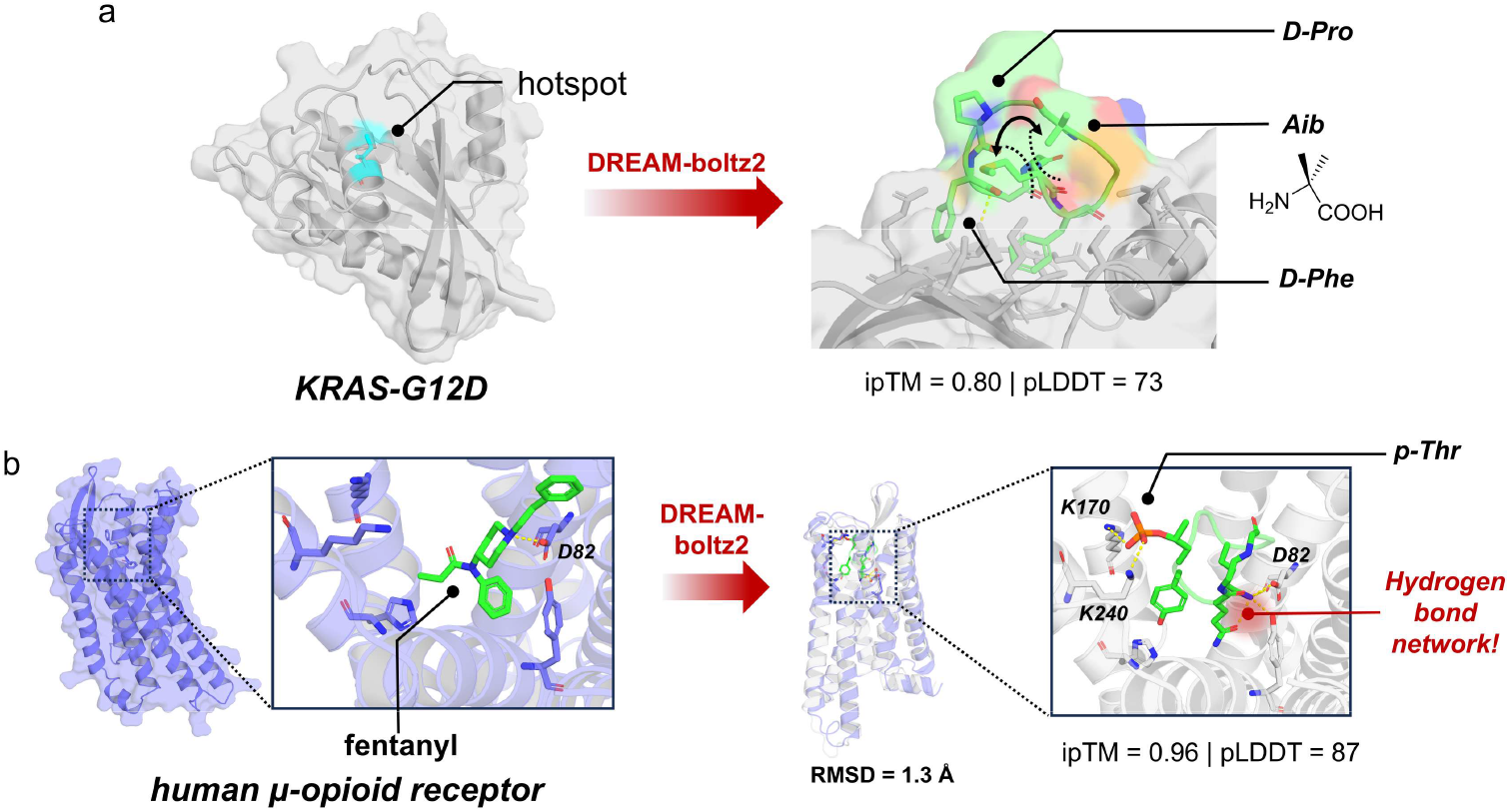
Programmable Therapeutics for Complex Targets. (a) Precision Scaffolding for KRAS-G12D: Targeting the shallow, “undruggable” surface of KRAS with a compact 10-mer cyclic peptide. The model orchestrated a non-canonical ensemble to achieve high-affinity binding: D-Proline and D-Phenylalanine induce a sharp reverse turn, allowing the peptide to execute a “pincer maneuver” into the hydrophobic cleft. Crucially, an Aib residue utilizes its gem-dimethyl group to exert “steric steering” against the receptor surface, locking the bioactive conformation. (b) Next-Generation Opioid Mimicry: Design of a peripherally restricted agonist for hMOR. Unlike the lipophilic small molecule Fentanyl (PDB: 8EF5)^25^ (left), which relies on deep hydrophobic burial, the DREAM-designed peptide (right) adopts an electrostatic binding logic. The model autonomously selected a Phosphothreonine (p-Thr) to bridge extracellular Lysines (K170, K240), creating a “vertical lock” that stabilizes the active conformation (RMSD = 1.3 Å). This spontaneous shift towards a highly polar, salt-bridge-dependent binding mode naturally aligns with the design goal of reducing blood-brain barrier permeability.

We next applied this framework to design a safer alternative to Fentanyl, trying to address the “Holy Grail” challenge for potent but non-addictive analgesia^26^. Traditional agonists like Fentanyl are highly lipophilic, enabling them to cross the blood-brain barrier (BBB) and trigger central side effects, including addiction and lethal respiratory depression^27^. Our objective was to design a cyclic peptide mimic that retains high potency while remaining peripherally restricted due to its modified molecular properties. Structural analysis again reveals that the model exhibited a level of emergent chemical intuition reminiscent of a seasoned medicinal chemist. Without prior instruction to prioritize polar residues, the optimizer autonomously identified two extracellular Lysines (K170 and K240) as high-energy electrostatic hotspots. To neutralize this positive cluster, the model bypassed canonical residues and specifically selected a Phosphothreonine (p-Thr) to serve as a high-density electrostatic anchor (Figure 3b). This selection achieves what we term a “vertical lock” mechanism: while the peptide’s core scaffolds into the classical deep-seated pocket, the p-Thr creates a sophisticated, multi-centered salt-bridge network at the receptor’s vestibule. By autonomously shifting from the lipophilic architecture typical of traditional opioids to this electrostatic-led binding logic, the design demonstrates the framework’s inherent ability to prioritize biophysical features — such as increased polar surface area—that naturally hinder membrane translocation. This suggests that DREAM-boltz2’s optimization process is sensitive to the intrinsic biophysics of the target interface, potentially offering a structural basis for addressing therapeutic challenges like systemic toxicity through de novo design.

## Discussion

Our work extends the “predict-to-design” paradigm—previously established for Evoformer-based models like AlphaFold2—to the new generation of all-atom diffusion predictors. While methods such as BindCraft have successfully inverted deterministic predictors, extending this capability to stochastic diffusion models has remained challenging due to the computational prohibitive cost and gradient instability associated with long sampling trajectories. DREAM bridges this gap via Geometric Score Distillation (GSD). By leveraging the single-step denoised estimate 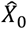 as a differentiable proxy, we validate that the generative potential of diffusion-based predictors can be unlocked without specialized retraining.

A pivotal observation from this inversion is the model’s access to a “latent chemical vocabulary.” The frozen Boltz-2 predictor exhibited an emergent capacity for sophisticated medicinal chemistry, autonomously deriving physical solutions rather than merely matching patterns. This was evident in the spontaneous selection of D-amino acids to resolve chiral clashes in B7-H3, the engineering of a figure-8 bicyclic scaffold to minimize entropy on PD-L1, and the deployment of Aib for steric locking in KRAS. Particularly in the case of hMOR, the selection of Phosphothreonine to create an electrostatic lock demonstrates a biophysical intuition aligning with complex pharmacological goals.

Looking forward, the “Energy-Anchored” strategy — fundamental to our framework’s acronym—outlines the path from topological scaffolding to atomic precision. While the current results demonstrate that high-noise regimes (σ ≈ 500) are sufficient for shaping global topologies and identifying hotspots, accessing fine-grained physicochemical features (e.g., specific hydrogen bond networks) requires mitigating the averaging effect of high-sigma estimation. As detailed in our Methodology, anchoring the gradient calculation to valid, low-energy conformations enables operation in low-noise regimes. This capability will be essential for future applications requiring sub-angstrom precision, such as nanobody design or catalytic site engineering, positioning DREAM as a scalable, model-agnostic engine for next-generation molecular discovery.

## Acknowledgements

We gratefully acknowledge the open-source community for the codebases used in this work, particularly Boltz and PyTorch. We thank BJMUHPC for providing high-performance computing resources. Additionally, Yuxuan Li would like to express special thanks to Jili*X*ie and Yihaofan Jiang for helpful discussions, and to Wei Ren for his assistance with wet-lab experiments.

